# Seasonal dynamics and repeatability of gastrointestinal parasite burdens in wild Soay sheep

**DOI:** 10.1101/2021.03.10.434794

**Authors:** Amy R. Sweeny, Yolanda Corripio-Miyar, Xavier Bal, Adam Hayward, Jill G. Pilkington, Josephine M. Pemberton, Tom N. McNeilly, Daniel H. Nussey, Fiona Kenyon

**Author notes:** Author for correspondence: Amy R. Sweeny.

## Abstract

Seasonality is a ubiquitous feature of wildlife disease ecology, but is determined by a complex interplay of environmental, parasitological and host factors. Gastrointestinal parasites often exhibit strong seasonal dynamics in wild vertebrate populations due to, for example, environmental influences on free-living or vectored life stages, and variation in the physiological and immune status of hosts across their annual cycle. At the same time, wild populations are typically infected with multiple parasites. The seasonal dynamics of co-infecting parasites may differ depending on age and reproductive status, and associations among parasites may be driven by short-term within-individual changes or longer-term interactions that are consistent among hosts. Here, we used faecal samples and egg counts collected repeatedly from individually marked and monitored wild Soay sheep that were part of a long-term study to investigate seasonal dynamics of six gastrointestinal parasite groups (strongyle nematodes, coccidian protozoa, *Capillaria, Strongyloides, Nematodirus*, and *Moniezia*). Prevalence and abundance generally tended to be higher spring and summer, and burdens were higher in lambs than adults. Within the highly prevalent strongyle nematode group, we found differences in seasonality of egg counts depending on adult reproductive status. Reproductive ewes had increased counts in spring around the time of birth followed by a drop in abundance in summer, while barren ewes showed little evidence of seasonality. Males showed a sustained rise in egg counts through spring and summer, and sex differences were only strongly apparent in summer. In contrast, in similarly prevalent coccidia we found a peak in faecal oocyst counts in spring but no differences in seasonality among males, barren and pregnant ewes. Using multivariate mixed-effects models, we went on to show that both strongyle and coccidia counts are moderately repeatable across seasons among individuals. We further show that apparent positive correlation between strongyle and coccidia counts was driven by short-term within-individual changes in both parasite burdens rather than long-term among-individual covariation. Overall, our results demonstrate that seasonality varies across demographic and parasite groups and highlight the value of investigating fluctuating susceptibility and exposure over time for understanding epidemiology of a population.

## 1. Introduction

Infection dynamics in wild animal populations are complex: variation in the environment will impact parasites to influence exposure and transmission, as well as host physiology to influence susceptibility and resistance (Acevedo-Whitehouse and Duffus, 2009; Cardon et al., 2011; Hawley and Altizer, 2011; Stromberg, 1997). These environmental influences incorporate short-term variation (e.g. seasonal cycles) and longer-term repeatable differences among hosts (e.g. host home range quality and early-life environment; (Altizer et al., 2006; Hayward et al., 2010; Smith et al., 1999). Furthermore, wild populations are typically infected with a diverse suite of parasites, each potentially showing varying sensitivity to environmental pressures due to differences in their life cycles and vulnerability to different types of host immune response (Cox, 2001; Pedersen and Fenton, 2007; Petney and Andrews, 1998). It is also well established that parasite burdens vary with host age and sex (i.e. demographic group), and the exposure and susceptibility of these different host groups may respond in different ways to variation in the environment (Hayward et al., 2009; Zuk and McKean, 1996). In order to better understand the complex parasite dynamics of natural systems, we need non-invasive, longitudinal studies of well-understood wild populations which monitor important parasite taxa repeatedly in individuals from different demographic groups (Clutton-Brock and Sheldon, 2010).Here, we use a long-term, individual-based study of wild Soay sheep (*Ovis aries*) to determine the seasonal dynamics and within-individual repeatability of a range of gastrointestinal (GI) parasites, and to test how demographic groups within the population respond differently to season variation.

Seasonality in parasite dynamics is commonly observed in wild vertebrates, but manifests differently depending on parasite and host population characteristics (Albery et al., 2018; Altizer et al., 2006; Martin et al., 2008; Nelson and Demas, 1996). Seasonal fluctuations in the weather and resource availability can act directly on parasites and their vectors, for instance by influencing survival of vectors or environmental stages of parasites, limiting or enhancing exposure (Altizer et al., 2006). Seasonal fluctuations can also impact the physiological state of hosts to impact their exposure, susceptibility and resistance to parasites. For example, reduced food availability in winter reduces the energetic resources available to hosts, which is expected to reduce investment in immunity (Martin et al., 2008; Nelson and Demas, 1996; Zuk and Stoehr, 2002). In seasonal breeders, the arrival of large numbers of highly susceptible, immunologically naïve juveniles into the population at a certain point in the year may raise transmission and exposure across the population as a whole (Wilson et al., 2004). Furthermore, host immunity fluctuates seasonally due to temporal variation in energetic demands and constraints as associated with growth and reproduction (Martin et al., 2008; Nelson and Demas, 1996). Juveniles may be particularly susceptible to infection and pathology due to their poorly developed immune response and diversion of limited resources away from immunity and into growth (Ashby and Bruns, 2018; Loveridge and Macdonald, 2000; van Dijk et al., 2013). Sex differences in parasite burden are often observed, particularly in polygynous species including many mammals, and explained in terms of differences in resource allocation (Moore and Wilson, 2002; Zuk and McKean, 1996). In such systems, males tend to have higher parasite burdens than females, and this is ascribed to the high costs of intrasexual competition for mates limiting investment in immunity in males (Ezenwa et al., 2012; Muehlenbein and Bribiescas, 2005; Zuk and McKean, 1996). However, there is also evidence that seasonal female investment in reproduction has costs which impact on immunity and parasite burden (Metcalf and Graham, 2018; Zuk and McKean, 1996). This is perhaps most notably in female mammals, which often show a peak in infections around birth (the so-called peri-parturient relaxation of immunity, PPRI) that is ascribed to reduced immunity during late gestation and early lactation (Ayalew and Gibbs, 2005; Brunsdon, 1970; Houdijk, 2008; Houdijk and Jessop, 2001). Thus, it is well-established that season, age and sex impact parasite dynamics across wild and managed vertebrate systems, and there is good reason to expect seasonal effects to differ among host demographic groups. However, studies to date have tended to study parasite seasonality within one demographic group or a single parasite group, and our understanding of the interactions between season, age and sex in driving parasite community dynamics in the wild remains limited.

When multiple parasites are endemic in a population, as is the norm in the wild, seasonal fluctuations can be idiosyncratic to specific species or groups within the wider parasite community. How parasites with different transmission and lifecycles respond to environmental conditions may underlie this variation (Cable et al., 2017). For example, a recent study of wild red deer showed striking differences in seasonality between environmentally transmitted GI strongyle nematode parasites, a tissue-dwelling helminth parasite, and an invertebrate-vectored liver fluke (Albery et al., 2018). Importantly, even among environmentally transmitted parasites inhabiting the same site within the host, we can observe differences in burdens among demographic group and season. For example, gastrointestinal (GI) strongyle parasite burdens in domestic and wild sheep are observed to peak around the breeding season in adult females (via the PPRI), but peak later in summer and autumn in lambs (Hamer et al., 2019; Vlassoff et al., 2001; Wilson et al., 2004). In contrast, parasitic GI apicomplexans of the genus *Eimeria* are found disproportionately in immature lambs and burdens across the population peak later in spring and summer aligned with increased transmission via the pasture after infections establish in young individuals (Chartier and Paraud, 2012). Furthermore, co-infecting parasites sharing host sites and resources can interact to powerfully impact infection dynamics (Ezenwa, 2016; Fenton, 2008; Graham, 2008). For instance, in wild wood mice removal of a key helminth species (*H. polygyrus*) resulted in a 15-fold increase in coccidia burdens via a hypothesised competitive release (Knowles et al., 2013; Rynkiewicz et al., 2015). Clearly, longitudinal studies monitoring environmental and demographic variation across different parasite groups within host populations are important for improving our understanding of complex infection dynamics in wildlife.

Alongside shorter-term environmental influences on infection dynamics, several studies of wild mammals have observed differences in the average parasite burdens of individual hosts which are persistent over longer stretches of time (Albery et al., 2018; Debeffe et al., 2016). Such repeatable differences may be due to genetically-based differences in immunity or behavior, and there is mounting evidence that parasite burdens and immune measures are heritable in wild vertebrates (Gasparini et al., 2009; Gold et al., 2019; Graham et al., 2010; Hayward et al., 2014; Råberg et al., 2003). Alongside such genetic effects, early-life environmental conditions can also shape lifelong differences in parasite burdens. For example, in European shags (*Phalocrocorax aristotelis*), environmental conditions during rearing had stronger impacts on chick nematode burdens than did host factors (Granroth-Wilding et al., 2014). Consistent among-host differences in space use are also likely to contribute to repeatable differences in infection rates (Albery et al., 2019; Debeffe et al., 2016; Shaw and Dobson, 1995) and there is some evidence that the repeatability of parasite burden may differ depending on the parasite in question in natural systems (Albery et al., 2018). Estimating the repeatability of parasite burdens and understanding its causes remains an important topic for wildlife disease ecology and host-parasite co-evolution for understanding influence of, e.g. genetic variance and response to selection. However, an important and rarely studied related question is whether associations among parasite species observed within a host reflect short-term dynamic changes in burdens or long-term repeatable differences among hosts in their parasite community composition. Recently, longitudinal data and hierarchical mixed-effects models have been used to test whether associations between parasite burdens and body condition and between immune measures and survival are driven by within- or among-individual level associations in wild mammals (Debeffe et al., 2016; Froy et al., 2019). However, to our knowledge, no study has applied this approach to dissect the within- and among-host drivers of correlations between co-infecting parasites, despite the potential significance of this for the dynamics of natural infections.

The Soay sheep living on the St Kilda archipelago represent a powerful system to disentangle the roles of long-term individual differences and short-term effects of environment, age and reproductive status on parasite dynamics in the wild. Individual sheep in the study population are uniquely marked at birth and subsequently closely monitored, and are repeatedly caught and sampled across their lifetimes. The parasite community of this population is well characterised, with environmentally transmitted GI nematodes and coccidians as the primary parasites (Graham et al., 2016; Wilson et al., 2004). Higher burdens of both parasite groups are expected in the spring and summer due to faecal-oral transmission cycles and the abundance of immunologically naïve lambs who harbour disproportionately high intensities of infection which increase exposure in the environment (Craig et al., 2007; Wilson et al., 2004). In addition, for strongyle nematodes, there is evidence that males tend to have greater parasite burdens than females, and reproductively active females exhibit a strong peak in the spring due to periparturient relaxation in immunity (PPRI) associated with reduced ability to respond to infection surrounding giving birth (Wilson et al., 2004). To date, however, longitudinal parasitological study in this system has been limited to spring and summer and has focused on the strongyle nematode group only. Here, we investigate seasonal dynamics of 6 GI parasite groups by repeatedly faecal sampling known individuals belonging to different age and sex groups across five sampling points within a single year. We test the predictions that: (i) parasite burdens will be highest in warmest months and (ii) in the youngest individuals; (iii) males will have higher burdens than females across the year, and (iv) parasite burdens will be raised in spring among breeding females due to PPRI. We specifically test whether seasonal dynamics vary among age and sex groups, as well as estimating the individual repeatability of strongyle nematode and coccidian burdens and testing whether correlations between burdens of these two highly prevalent parasite groups are driven by among- or within-individual level processes.

## 2. Methods

### 2.1. Data collection & study system

This study was conducted in a free-living population of Soay sheep on the St Kilda archipelago (57°49’N, 08°34’W, 65 km NW of the Outer Hebrides, Scotland). A population of 107 Soay sheep was moved from the island of Soay to the largest island of the archipelago, Hirta, in 1932. The resultant unmanaged Soay sheep population living in Village Bay on Hirta has been part of a long-term individual-based study since 1985 (Clutton-Brock et al. 2004). The annual reproductive cycle of the population is illustrated in Figure 1. Fieldwork to monitor the population and collect samples and data follows a similar pattern. In April, most ewes give birth to one or two lambs (approximately 20% of births are twins), and neonates are captured within a few days of birth and marked with unique ear tags allowing them to be identified and monitored throughout life. Sheep gain condition and females lactate to their lambs over the months that follow, with most lambs becoming largely weaned by four months old. In August, as many individuals resident in the Village Bay population are caught in corral traps as possible (usually 50-60%) over a two week period. Morphological measures are taken, along with blood and faecal samples, at capture. Males live in bachelor groups for much of the year and gain body condition through spring and summer in preparation for the autumn rut, during which males compete for mating opportunities with oestrous females. A fieldwork team monitors the rut and captures and marks incoming, immigrant males in October and November. In the winter months that follow, the sheep are food limited and experience challenging climate conditions, and most natural mortality occurs late in the winter period (February and March). The majority of carcasses of study animals are recovered during winter mortality searches so death can be determined with certainty in most cases.

**Figure 1.**
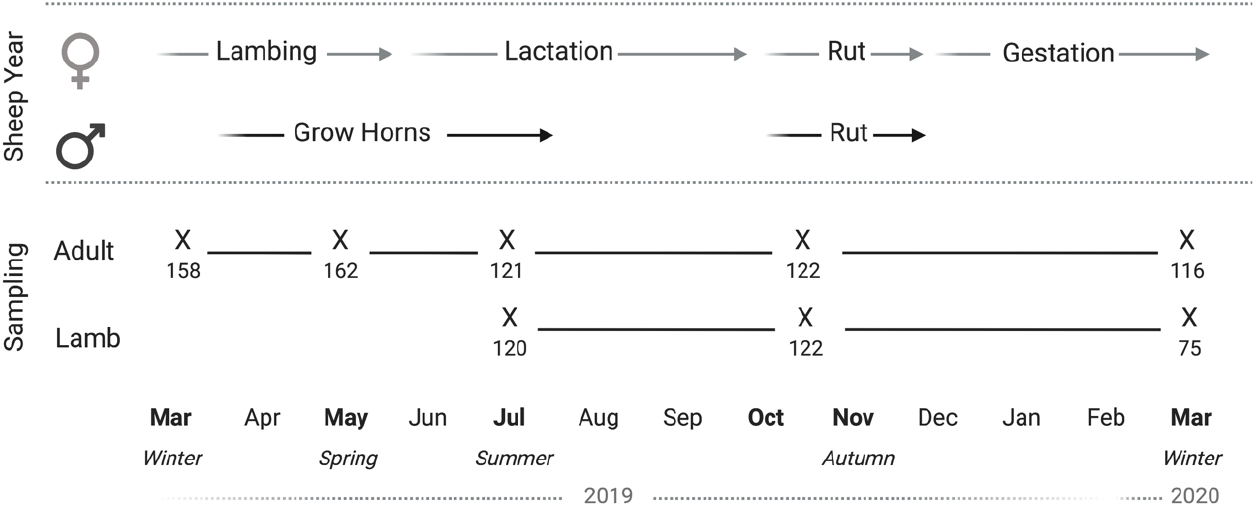
The study’s faecal sampling program (“sampling”) is illustrated alongside the major events of the annual cycle of Soay sheep on St Kilda (“sheep year”). Each ‘X’ designates each sampling point for adults or lambs with the number of samples collected indicated below each sampling point. Wherever possible, the same individuals as initially selected were repeatedly sampled across the study period.

Faecal samples for parasitological analyses were collected seasonally in five sampling trips to capture major seasonal variation: winter 2019 (6 March −16 March 2019), spring 2019 (13 May – 24 May 2019), summer 2019 (17 July – 29 July 2019), autumn 2019 (27 October −8 November 2019), and winter 2020 (4 March – 18 March 2020) (Figure 1). We targeted individuals from two demographic groups for repeated sampling over this period: adults (aged 2 years or more, sampled from winter 2019 onwards) and lambs (sampled from summer 2019 onwards). Lambs were not sampled in spring 2019 despite being alive then because faecal samples are used for multiple purposes within the broader study system and their small size meant they would not consistently produce a sample of sufficient size; furthermore earliest non-zero FECs from lambs are from June onwards (Wilson et al., 2004). Selected individuals included as even a distribution of males and females as possible, given the adult population is heavily female-biased (Wilson et al., 2004). Target individuals were observed closely until they defecated. The faecal sample produced was collected from the pasture within 2 minutes of defecation. For each sample, individual identity, collection date and time of day was recorded. Samples were weighed at the end of each collection day. 2g of faecal matter for adults and 1g for lambs were stored anaerobically in bags to minimise oxygen exposure. Samples were refrigerated at 4°C until processing to minimise egg hatching. Parasite counts were carried out upon return to the laboratory within 3 weeks after collection.

### 2.2. Parasitology

Gastrointestinal parasites were quantified using a modified salt-flotation method to enumerate eggs (helminths) or oocysts (protozoans) in the faeces (Hayward et al., 2019). Briefly 2g (adults) or 1g (lambs) of faecal matter was mixed with 10 mL of water per gram and mechanically homogenised. A 10-mL aliquot of each sample suspension was filtered through a 1mm sieve and washed with 5-mL of tap water. Filtrates were centrifuged in 15-mL polyallomer tubes for 2 min at 200 g. The supernatant was removed and faecal pellet resuspended with 10mL of saturated NaCl solution and centrifuged for another round of 2’ at 200 g. Tubes were clamped below the meniscus using medical forceps, leaving gastrointestinal parasites eggs or oocysts which float in salt solution in the fluid above the clamp. Fluid above the clamp was added to a cuvette in addition to approx. 1mL of NaCl used to wash the upper chamber. The cuvette was topped up with NaCl solution and the entire cuvette surface scanned to count parasite eggs and oocysts to a precision of 1 egg/ occyst per gram (EPG/OPG). Parasites identified in counts included strongyles (primarily *Teladorsagia circumcincta* and *Trichostrongylus spp*., as well as *Chabertia ovina* and *Bunosomum trigonocephalum*), coccidia (*Eimeria spp*.), *Strongyloides papillosus, Capillaria longipes, Nematodirus spp*., and *Trichuris ovis*. Segments of the tapeworm *Monezia expansa* were also identified via salt flotation method, and scored as presence or absence. We use the term abundance to describe the faecal egg/oocyst counts (FEC/FOC) for strongyles, coccida, *C. longipes*, and *Nematodirus spp*. We use presence/absence scores to describe infection probability for *S. papillosus, M. expansa*, and *T. ovis*, which were much rarer in the population. FECs are a common proxy for worm burden in wild animals and represent a good approximation of burden in Soay sheep (Wilson et al. 2003), and high repeatability with the cuvette method described above has previously been established in Soay sheep (Hayward et al., 2019).

### 2.3. Statistical analysis

All statistical analyses were conducted using R version 3.6.1 (R Core Team 2019). We fitted all models using the Bayesian modelling package ‘MCMCglmm’ (Hadfield, 2010). All models detailed were run for 260,000 iterations with a 2,000-iteration thinning interval and a 60,000-iteration burn-in period. We used generalised linear mixed-effects models (GLMMs) to test the effects of season and sex on egg/oocyst counts for each parasite group separately, and ran separate models for lambs and adults due to differences in the seasons sampled in each age group (Figure 1, Table 1). *Trichuris* had a prevalence of <1% in the population (Figure 2) and was not included in our analyses. *Capillaria* was found only in adults and therefore does not appear in lamb analysis, and *Nematodirus* were found almost exclusively in lambs and therefore does not appear in adult analysis (Figure 2). In few instances, parasites had zero or close-to-zero prevalence for some time points so models were fit to subsets of the data reflecting this (Figure 2; Table 1). Strongyle, coccidian, *Capillaria*, and *Nematodirus* models were fitted with FEC/FOC as a response variable and Poisson error families. *Strongyloides* and *Moniezia* were scored as presence/absence and therefore modelled with a binomial error family (Table 1). Individual identity was included as a random effect in all models. We assessed statistical support of differences between levels of season for each parasite by comparing the proportion overlap of the posterior distributions of the MCMC estimates for each level divided by half the number of stored iterations (Albery et al., 2018; Palmer et al., 2018).

**Figure 2.**
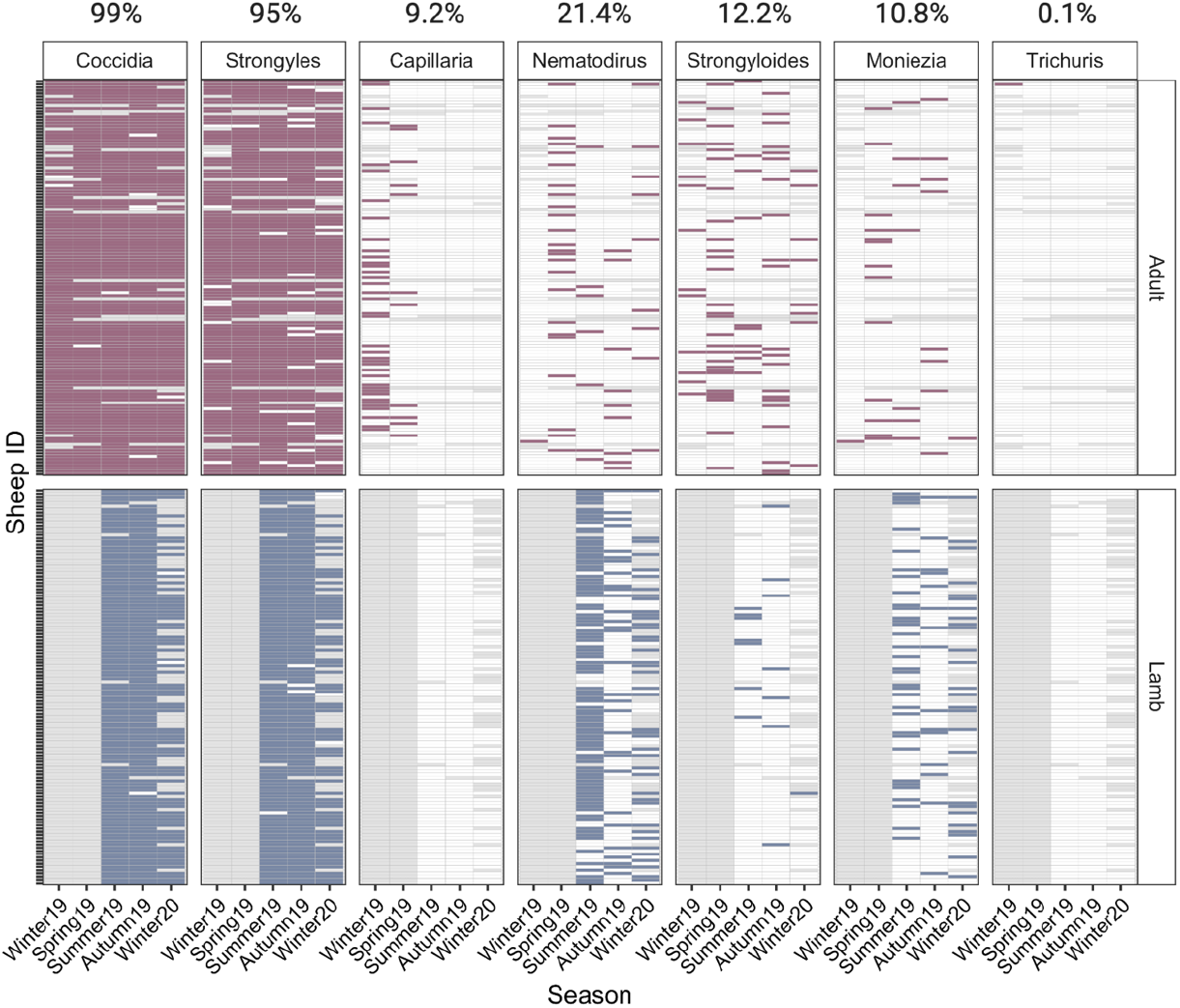
Infection status of individual sheep across parasite groups and seasons. Shading represents the presence of a parasite for each sample (filled: present; blank: absent; grey: not sampled). Top panel represents adult samples and bottom panel represents lamb samples. Overall prevalence within the population is shown for each parasite at the top of the plot.

**Table 1.**
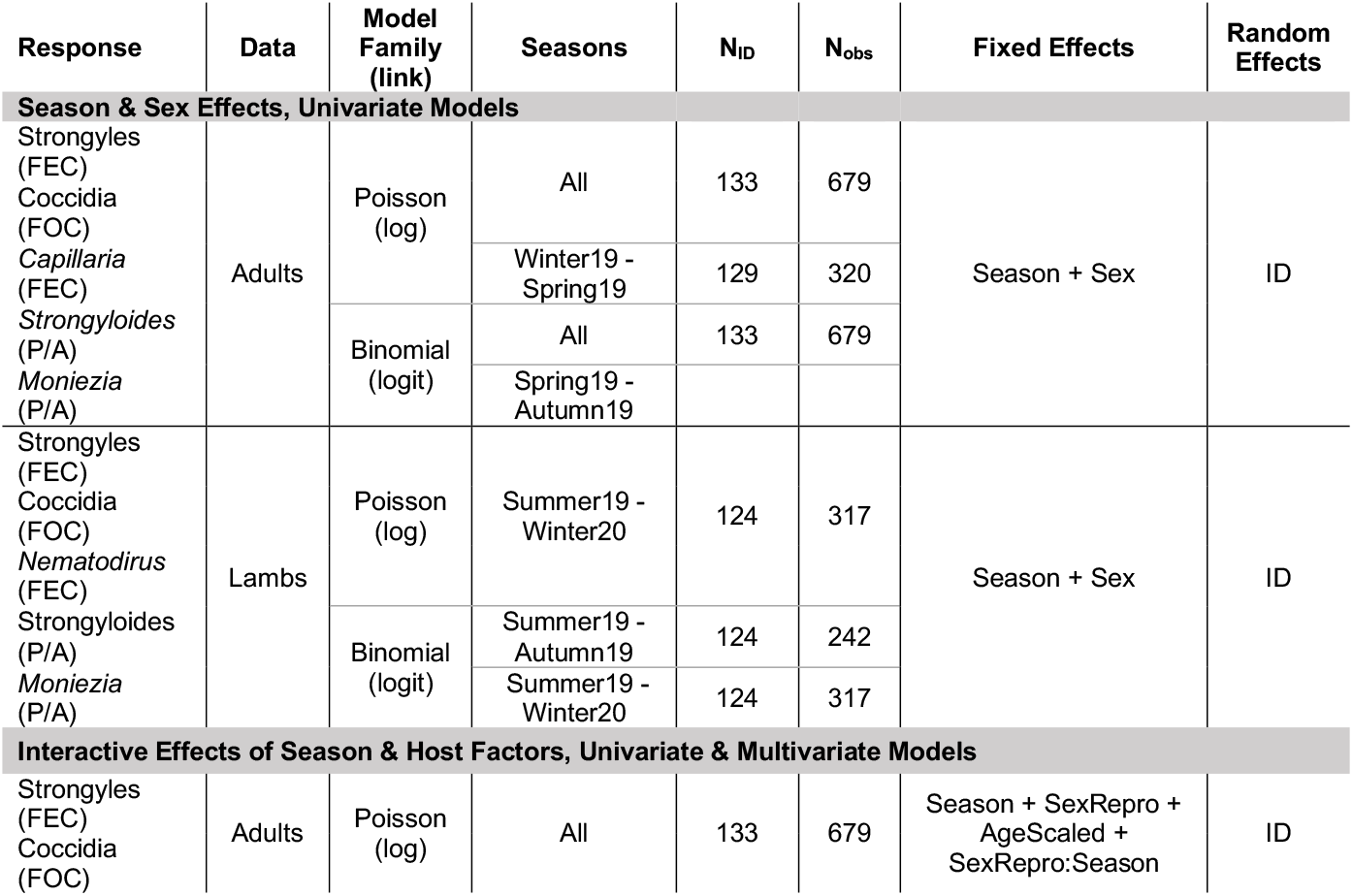
Details of generalised linear mixed-effects models fit to parasite data in this study (see text for further details).

We then investigated whether and how observed seasonal dynamics in parasite counts were shaped by interactive effects with sex and reproductive status in adult sheep. We limited these analyses to strongyle egg and coccidian oocyst counts only, as these were the two most prevalent parasites in the population and other parasites had very low prevalence in some seasons (Figure 2). To test for sex differences overall as well as differences among females that did or did not breed, we coded a ‘sex / reproductive status’ variable with the following categories: “male”, “female no lamb” (did not breed in spring 2019) and “female with lamb” (gave birth to at least one lamb in spring 2019). GLMMs with Poisson error families were fitted to both strongyle FEC and coccidia FOC with the following fixed effects: sex / reproductive status (“SexRepro”), Season (5 level factor, as above), Age in years (continuous, scaled to a mean of zero and a standard deviation of 1), and a SexRepro-by-Season interaction (Table 1). Individual identity was included as a random effect in these models. We used posterior prediction to confirm model performance against raw data, and to investigate whether and how SexRepro categories varied in their seasonal trajectories. 1000 predicted values were generated for each SexRepro/Season category to calculate a distribution (Season_t+1_-Seasont) representing the relative change from season-to-season across sampling period. Resultant distributions were compared across each combination of SexRepro categories for each seasonal change to assess significant differences in seasonality for demographic groups, where significance values were calculated using proportional overlap of each pairwise set of distributions divided by half the number of predicted values (equivalent to a post-hoc test).

Finally, we investigated how repeatable strongyle and coccidia counts were for adults across sampling seasons, as well as the among- and within-host covariance in strongyle and coccidia counts. We specified a multivariate Poisson model with strongyle and coccidia counts as response variables, and fixed and random effects identical to those in univariate models above (Table 1). Individual repeatability across seasons for both responses was derived using the individual variance estimate from each model divided by the total variance (Var_ID_ / (Var_ID_ + Var_Residual_ + Var_Poisson_) (Nakagawa and Schielzeth, 2010). This model also estimates the covariance at the among-individual and within-individual (residual) level between strongyle FEC and coccidia FOC. The covariance at the among-individual level (Cov_Individiual_) indicates the association between individuals’ average strongyle FEC and coccidia FOC across the entire study period. The within-individual or residual covariance (Cov_Residual_) reflects associations at the level of sample point having accounted for any association at the individual mean level. Overall (phenotypic) covariance between the two responses is then represented by Cov_phenotypic_ = Cov_Individual_ + Cov_residual_.

## 3. Results

Strongyles (95% prevalence) and coccidia (99%) were by far the most prevalent parasites in the population, followed by *Nematodirus* (21.4%)*, Strongyloides* (12.2%), *Moniezia* (10.8%) and *Capillaria* (9.2%; Figure 2). For parasites with associated count data (versus presence/absence), mean and range of counts for each age and season group is summarised in Table S1. There was seasonal variation in prevalence of all parasites (Figures 2). coccidia and strongyles were present at high prevalence in both adults and lambs for all seasons sampled, with the exception of a decrease in prevalence for lambs during winter 2020. In contrast, other parasites were much less consistent in detection. *Capillaria* was found only in adults in the first two seasons and then not again. *Nematodirus* was found primarily in lambs especially in the summer, but infrequently in adults across all seasons as well. *Strongyloides* was present primarily in adults and detected most frequently in spring and summer, while *Moniezia* was detected in low overall prevalence but more frequently in lambs. Finally, *Trichuris* was detected in only one adult and was dropped from further analysis.

Our models of parasite counts likewise revealed seasonal variation for most parasite species groups (Figure 3). In adults, parasite abundance was generally significantly higher in spring than in other seasons and, while it tended to be lower in summer than spring, summer counts tended to be higher than autumn or winter seasons (Figure 3A-E, Tables S1–2). For strongyle and coccidian counts, there was a clear seasonal pattern where spring FEC/FOCs were greater than summer, and both spring and summer were greater than autumn and winters, which were more similar in magnitude (Figure 3A & B, Table S1). *Capillaria* was only detected in winter and spring 2019 and had highest abundance in the winter (Figure 3C). Probability of detection was highest in spring for *Strongyloides*, although seasonal patterns for this parasite were less consistent than strongyles and coccidia (Figure 3D). There was no difference in probability of infection with Moniezia across seasons (Figure 3E). In addition to seasonal effects, we found that males had higher strongyle egg counts than females (post. mean = 0.97, 95% CI: 0.5, 1.45, P_MCMC_ < 0.001) and males had greater probability of infection with *Strongyloides* than females (post. mean = 1.28, 95% CI, 0.21, 2.38), P_MCMC_ = 0.03). There were no sex differences in the other parasite groups for adults (Table S2). Although parasites were modelled within age groups, parasite abundance was generally higher in lambs than in adults save for *Capillaria* (Table S1).

**Figure 3.**
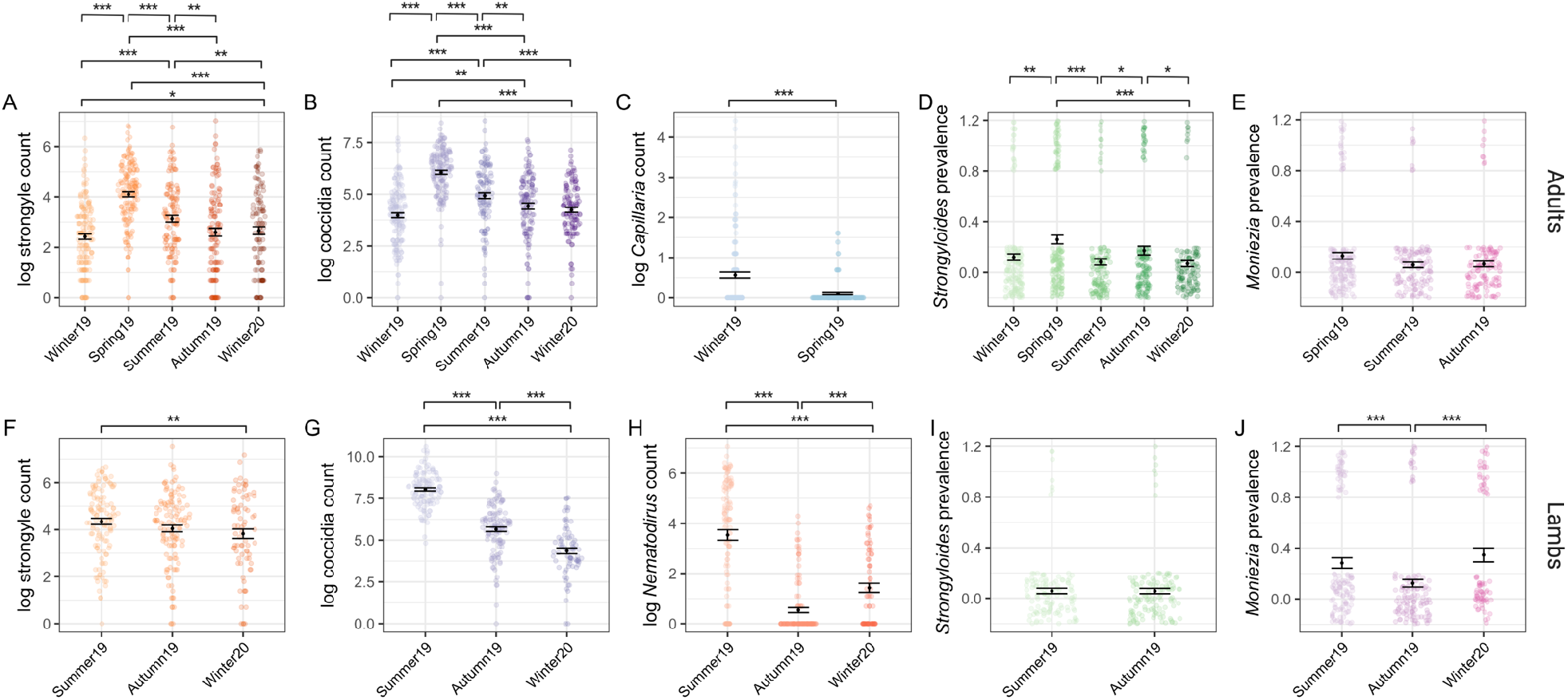
Seasonal dynamics of six gastrointestinal taxa of Soay sheep spanning five sample trips. Top panel (A-E) represents data from adults (all sampling points). Bottom panel (F-J) represents data from lambs (Summer 2019 onwards). Coloured points represent raw data, where the width of the point spread is proportional to the density distribution. Black points and error bars represent the mean (for counts) or prevalence (for presence/absence) for each parasite ± SE. Brackets above indicate pairwise comparisons across all season combinations, where significance was calculated using as the proportional overlap between posterior distributions for pairs of Season levels divided by half the number of stored iterations. Significance is denoted by ***, ** and * for P<0.001, P<0.01 and P<0.05 respectively. Some models were run in adults (*Capillaria*) or lambs (*Nematodirus*) or only run on subsets of data due to zero-extremely low prevalence within some time points. Plots and effect comparisons are limited to those categories represented in the models.

In lambs – for which we only had samples in summer and autumn 2019 and winter 2020 – we found that strongyle and *Nematodirus* FECs and coccidian FOCs were highest in summer (Figure 3F-H; Table S2). The probability of detecting *Moniezia* eggs was lower in autumn than either summer or winter (Figure 3J), whilst there was no difference in the probability of detection of *Strongyloides* eggs between summer and autumn and this parasite was detected in only one lamb in winter 2020 (Figure 3I; Table S2), We found no evidence for differences between male and female lambs in their average counts across seasons in any of these parasite groups (Table S2).

Models investigating interactive effects of sex / female reproductive status and season demonstrated much stronger interactions for strongyle FECs than in coccidian FOCs (Figure 4; Tables S3–4). Strongyle counts increased significantly more from winter to spring in females that produced lambs than in males (Figure 4 A & C). Females that did not produce a lamb in contrast had a significantly lower magnitude of change from winter to spring and these barren ewes showed little seasonal variation in FEC overall (Figure 4A & C). From spring to summer, strongyle FECs dropped in females with lambs to levels similar to those observed in females without lambs (Figure 4A & C), but male FECs remained high and actually increased slightly on average from spring to summer (Figure 4A & C). Coccidian counts showed little evidence of a season-by-sex/reproductive status interaction in contrast: counts increased from winter to spring in a consistent manner across sex / reproductive status groups, and then fell from spring to summer, although this drop was less pronounced in males than in females (Figure 4B & D). In both models, continuous age of adults was fit to account for potential ageing effects within the adult age class, but was insignificant for both strongyles and coccidia (Table S3).

**Figure 4.**
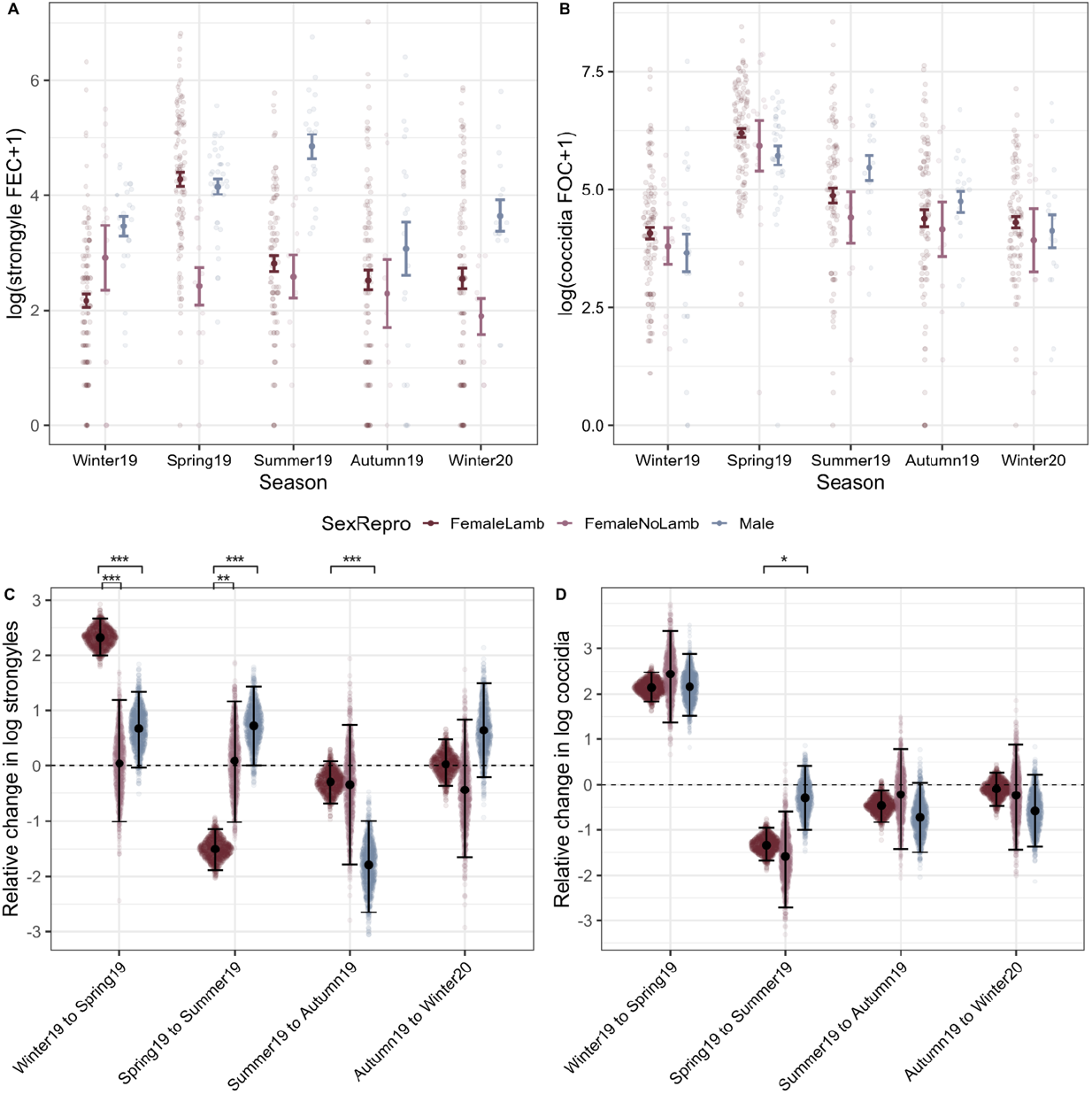
Interactions with season and host sex and reproductive status in adult sheep. Top panel (A-B) represents raw data (coloured points) overlaid with mean and SE grouped according to SexRepro-by-Season groups for both (A) Strongyle egg and (B) Coccidia oocyst counts. Maroon indicates females who gave birth to at least one lamb in spring, pink is females who did not give birth and blue is males. Bottom panel (C-D) represents the pair-wise comparisons of seasonal change calculated from 1000 predicted values for each SexRepro-by-Season group from GLMMs with (C) Strongyle and (D) Coccidia counts as a response. Sina plots of points represent each value from the distribution (Seasont+1-Seasont) for each group, overlaid with mean and 95% confidence intervals. Significant difference between SexRepro groups is indicated by brackets above plots indicating where significance was calculated using as the proportional overlap of each pairwise set of distributions. Comparisons can be interpreted similarly to the following examples: panel C indicates that females with lambs show a much greater increase in FEC from winter to spring than both females without lambs and males, and panel D shows that females with lambs show a greater decrease from spring to summer compared to males. Significant differences between effects are indicates by ***, ** and * for P<0.001, P<0.01 and P<0.05 respectively.

Bivariate GLMMs yielded nearly identical fixed effects estimates to the univariate models described above (Table S3). Repeatabilities of strongyle FECs and coccidia FOCs across seasons, after accounting for effects of sex and season were 0.28 (95% CI: 0.19-0.36) and 0.20 (0.12-0.27), respectively. The overall phenotypic correlation between strongyle and coccidia counts was estimated as 0.27 (0.20 – 0.37, P_MCMC_ < 0.001). Underlying this, we estimated a higher positive within-individual correlation (post.mean = 0.46, 95 % CI: 0.31, 0.62) and a much weaker among-individual correlation which had 95% credible intervals which overlapped zero (post.mean = 0.15, 95 % CI: −0.03, 0.31).

## 4. Discussion

We found that most parasite groups varied in prevalence and abundance with season and, as predicted, these tended to by highest in the spring and summer. Strongyle nematodes and coccidia were highly prevalent across the entire study period in both lambs and adults, while other parasite groups were detected infrequently or only in certain age groups or seasons. These results largely agree with previous work in this system (Craig et al., 2007; 2006; Wilson et al., 2004), as well as in domestic sheep (Evans et al., 2021; Hamer et al., 2019). GI parasite communities differed between lambs and adults: with *Capillaria* and *Strongyloides* considerably more prevalent in adults and *Nematodirus* more prevalent in lambs (Figure 2). Strongyle FEC and coccidia FOC were generally higher in lambs than adults across seasons (Figure 3; Table S1). Although we predicted that males would have higher parasite burdens than females, following life history theory and previous comparative studies (Moore and Wilson, 2002; Zuk and McKean, 1996), we only detected male bias across seasons in two out of the six parasites studied (strongyles and *Strongyloides*). Our in-depth analyses of the highly prevalent strongyle and coccidia groups in adults revealed striking interactions between season and reproductive group in the former but not the latter. We also demonstrated that burdens of both groups are moderately repeatable across our year-long study period, and that counts are positively correlated overall. Bivariate modelling of these two parasite groups further revealed that this correlation was driven by short-term fluctuations in same direction across seasons, rather than consistent differences in FEC and FOC between individual sheep on average across the study period

The seasonal epidemiology of the GI nematode parasites investigated here is well studied and understood from veterinary work on domestic sheep (Vlassoff et al., 2001) and these dynamics seem remarkably robust to management practices (Hamer et al., 2019), as well as being consistent with previous findings in the unmanaged population of Soay sheep on St Kilda (Wilson et al., 2004). The seasonal pattern is broadly as follows: the widely observed increase in nematode FECs observed through spring is driven by the ingestion of over-wintering stage 3 larval (L3) nematodes from pasture by adult sheep. These larvae then develop to maturity inside the host and the PPRI in ewes around lambing results in a surge in egg shedding onto pasture. Immunologically naïve lambs are highly susceptible to nematode infection and their exposure and burdens increase rapidly over spring and summer due to both shedding from ewes experiencing PPRI and from the growing number of lambs (Vlassoff et al., 2001). An increase in egg shedding by pregnant ewes through spring is well established in domestic sheep (Vlassoff et al., 2001) and has also been observed in the Soay sheep on St Kilda (Leivesley et al., 2019). In domestic lambs, FECs peak in summer and autumn as more lambs become infected and shed eggs onto the pasture, after which egg shedding declines into winter as fewer eggs develop on pasture due to declining temperatures (Vlassoff et al., 2001). Previous work on St Kilda shows that counts of strongyle L3 on pasture follow this expected pattern, rising through spring to a peak around July/August after which they decline rapidly (Wilson et al., 2004). Strongyle eggs first appear in the faeces of lambs at around 45 days old, and small-scale longitudinal studies of lambs suggest that while lamb FECs rise to a peak around August, they subsequently remain high and may continue to rise again through late winter (Feb – April; (Wilson et al., 2004)). Although our sampling regime prohibited us from detecting the well-established increase in lamb FEC from spring to summer, our data confirm that, on St Kilda, lamb FECs remain high and do not drop as markedly through autumn and winter as in adults (Figure 3A & F). The reasons for this interesting difference between the early-life dynamics in unmanaged Soay sheep and domestic sheep remain to be determined, and certainly warrant further investigation. One possibility is that on St Kilda lambs FECs remain high due to the food limitation and high adult worm burdens experienced by Soay lambs during winter (Wilson et al 2004; Craig et al 2006), which may be largely alleviated by regular anthelmintic treatment and food supplementation in domestic sheep.

We found pronounced increases in strongyle FEC from the end of winter (March) to spring (May) among females that produced lambs, but not in barren ewes (Figure 4A). This supports previous observations of reproduction-induced increases in FEC in Soay sheep (Hayward et al., 2019; Leivesley et al., 2019) as well as numerous other wild animals such as bighorn sheep *Ovis canadensis* (Festa-Bianchet, 1989), red deer (Albery et al., 2020), flying foxes (Plowright et al., 2008), and birds (Knowles et al., 2009; Nordling et al., 1998) and is a well-defined phenomenon in domesticated sheep flocks (Hamer et al., 2019; Vlassoff et al., 2001). Increased counts in reproductive ewes can be attributable to both relaxed immunity (PPRI) due to the demands of lactation and gestation (Festa-Bianchet, 1989; Sheldon and Verhulst, 1996) or due to higher exposure from increased foraging behaviour to compensate for resource demands (Hutchings et al., 2002). Previous evidence documenting decreased strongyle-specific IgG antibody levels during the lambing season in Soay ewes suggests the former mechanism is at least partially responsible the FEC increase in reproductive ewes in this system (Hayward et al., 2019). The data from our study provide some further support for an immune-mediated explanation, as the FECs of reproductive ewes fall rapidly to levels observed in barren ewes by summer (July; Figure 4A). Breeding ewes are expected to experience high energetic demands through both spring and summer due to lactation, and therefore if increased exposure through altered foraging behavior was responsible for raised egg shedding we would expect reproductive females to have higher FEC than barren females through until the end of lactation in summer.

Previous studies, both in domestic sheep and the unmanaged Soays on St Kilda, have tended to overlook seasonal parasite dynamics in adult males, and our results demonstrate that male FECs show a seasonal pattern distinct from that seen in lambs, pregnant or barren ewes. Adult males show a sustained rise in strongyle FEC through spring and summer, unlike reproductive ewes which peak in spring and drop in summer (Figure 4A). Contrary to predictions for a persistent male-bias in parasite burdens across ages and seasons, we found raised strongyle FECs in adult males relative to reproductive females only in summer. This finding highlights the potential, and often overlooked, importance of males in the transmission dynamics of helminths in wildlife (Ferrari et al., 2004; Grear et al., 2009). Although further investigation is required to understand the causes of this sex difference, it is likely that some combination of differences in resource allocation trade-offs and exposure to pasture larvae in adult males compared to other demographic groups is responsible. Adult males invest their limited nutritional resources in horn growth and gaining weight and condition through spring and summer ahead of the autumn rut. The trade-off between testosterone production, investment in secondary sexual characteristics, and immunity is hypothesised to predispose males to higher susceptibility to infection via an immunocompetence handicap (Ezenwa et al., 2012; Nunn et al., 2009; Zuk and McKean, 1996), and we might expect this to manifest as raised male parasite burdens during the peak in exposure to strongyle larvae on pasture in July and August. In the St Kilda population, adult males tend to segregate spatially and often spend time in ‘bachelor groups’ outside of the autumn rut (TCB/Pemberton book). This spatial and social segregation of adult males through spring and summer may reinforce immunologically-mediate sex differences, as these heavily infected males may be more likely to expose each other to larvae on shared grazing ranging. We note high strongyle FEC in adult males during the rut has been associated with reduced time spent engaged in sexual activity, suggesting that the summer peak in male burdens may have fitness costs and impact sexual selection in this system (Wilson et al. 2004). Further work addressing the seasonal dynamics of mucosal immunity in adult males (as has been done for females recently on St Kilda, see Hayward et al 2019), the relationship between home range sharing and parasite burdens, and the effects of autumn burdens on rut performance and reproductive success in males will help understand the causes and consequences of the male summer peak in strongyle worm burdens.

Coccidian parasites showed a seasonal pattern broadly consistent with expectations: a marked peak in spring in adults followed by a decline from summer on, and substantially higher burdens in lambs than adults across seasons with a decline in lamb burdens evidence from summer to winter. However, in contrast to the patterns discussed above for strongyles, coccidian FOCs showed largely consistent seasonal dynamics across reproductive ewes, barren ewes and males (Figure 4B). It seems that reproductive status and investment impact seasonal variation in burdens in quite different ways across these two groups of GI parasites. The main driver of seasonal variation in coccidia in domestic sheep is thought to be pasture oocyst contamination by immunologically naïve lambs through spring (Chartier and Paraud, 2012). Although we did not have lamb samples from May to confirm this in the present study, previous work on the Soay sheep supports the central role of lambs in transmission and shows that coccidia FOCs are on average an order of magnitude greater in lambs than adults in summer (Craig et al 2007). Sheep develop immunity to coccidia over their first year of life, a response largely thought to be mediated by T helper type 1 (Th1) responses, and are thought to be largely resistant to pathology caused by these parasites after this point (Craig et al., 2007). One possible explanation for the lack of a PPRI effect or sex differences in coccidia FOCs, in contrast to strongyle FECs, may lie in differences in the way the immune response involved in controlling these parasite groups develops and trades off with investment in reproduction. Previous studies of mammals have observed that different parasite groups may be differentially impacted by reproduction (Albery et al., 2020; Becker et al., 2018; Rödel et al., 2016). Furthermore, resistance to helminths is dependent on T helper 2 (Th2) responses, which can be constrained by immune commitment to the Th1 responses required for resistance to microparasites such as coccidia (Graham 2008). We recently showed in wild Soay sheep that Th1-associated immune phenotypes predicted reduced coccidia FOCs, whilst Th2-associated phenotypes predicted reduced strongyle FECs (Corripio-Miyar et al, in prep). While this supports the role of functionally distinct aspects of the immune response in controlling these two different parasite groups, we also found that sheep mounting strong Th1 responses also tended to mount strong Th2 responses (Corripio-Miyar et al, in prep). It is possible that the immune responses that develop in sheep to coccidia during the first year of life are more effective or efficient than those against strongyles, and that the resource costs of mounting anti-coccidian responses are lower than those for strongyles and thus show less evidence of being influenced by reproductive status. However, further work testing the relationship between reproductive investment, T helper immunity and parasite burdens would be required to test this hypothesis.

Strongyle FECs and coccidia FOCs were both moderately repeatable at the individual level over our study period. This shows that variation in parasite burdens across a year are driven, at least in part, by consistent among-host differences in some aspect of genotype or environment. Previous estimates in our study system put the heritability of August FEC and FOC measures at 10-20% (Beraldi et al., 2006), which suggests at least some of the among-individual variation observed in the present study may have a genetic basis. Whilst previous experimental studies in wild rodents have identified negative interactions between GI coccidia and strongyles, which appear linked to competition over shared host resources and competitive release (Knowles et al., 2013; Rynkiewicz et al., 2015), our observational data confirm previous reports of a positive correlation between burdens with these two parasite groups in Soay sheep (Craig et al 2008 Parasitology). One plausible explanation, which also applies to observed positive correlations between Th1 and Th2 immune phenotypes in this population (Corripio-Miyar et al, in prep), is variation in resource acquisition and joint exposure to multiple parasites which is likely very common in natural systems. Sheep acquiring less resources, due to competitive ability or home range quality, may be less able to mount immune response to multiple parasites and may need to forage more widely and less selectively and risk greater exposure to multiple parasites. Here, we have utilized multivariate mixed-effects to dissect whether this correlation is driven by among- or within-host processes. Although rarely used to date in this context, previous studies have used this approach to dissect the role of among- and within-host processes in linking nematode parasite burdens in wild horses (Debeffe et al., 2016) and the role of among-host and among-site processes driving amphibian infection dynamics (Stutz et al., 2017). Our analyses demonstrate that the positive covariance between strongyle FEC and coccidia FOC is predominantly a within-individual correlation. In other words, it results from short-term (season to season) deviations from an individual’s mean parasite burden in the same direction for both types of parasite, and not from some individuals having consistently higher or lower mean FEC and FOC. This highlights the utility of this approach for our understanding the drivers of associations among parasites within communities, and suggests that short-term environmental fluctuations impacting host resource availability – rather than long-term genetically or environmentally driven among-host differences – are responsible for the observed correlation between strongyles and coccidia.

Overall, our results demonstrate that seasonality varies across demographic and parasite groups and highlight the value of investigating fluctuating susceptibility and exposure over time for understanding epidemiology of a population. Although we focus here on strongyles and coccidia at the group level, there is considerable diversity of species within both on St Kilda (Wilson et al., 2004). Recent advances in characterising the gastrointestinal community with metabarcoding methods using noninvasive faecal samples for both nematodes and coccidia (Avramenko et al., 2015; Evans et al., 2021; Vermeulen et al., 2016) can offer higher resolution understanding of GI parasite community dynamics. Additionally, integration of multiple immune markers can provide further insights regarding how multiple components of reproductive effort influence different components of immunity. This will be a crucial link in understanding dynamic stressors affecting immunity and the outcome across multiple parasites in coinfected systems. Finally, results here using just one year of sampling demonstrate the value of repeated observations at the individual level for parsing among- and within-individual contributions to variance. From an epidemiological perspective, long-term individual sampling can inform processes underlying complex dynamics of multiparasite systems.

## Author Contributions

All authors contributed to the development and planning of the study. XB and JGP collected samples. YCM and FK carried out parasitology. ARS analysed the data and led the writing of the manuscript with assistance from DHN and FK. All authors contributed critically to manuscript drafts and analysis.

## Acknowledgements

This work was funded by a large NERC grant (NE/R016801/1), and the long-term study on St Kilda was funded principally by responsive mode grants from NERC. We thank the National Trust for Scotland for support of our work on St Kilda, and QinetiQ and Kilda Cruises for logistical support. We thank the Ecology Within Team for input in the analysis and manuscript and Greg Albery for comments on manuscript draft. Figure 1 was created with Biorender.com.

## Electronic Supplementary Material

**Table S1.** Raw means and range of counts within the population

| Age Class | Season | Strongyles |  | Coccidia |  | Capillaria |  | Nematodirus |  |
| --- | --- | --- | --- | --- | --- | --- | --- | --- | --- |
|  |  | Mean | Range | Mean | Range | Mean | Range | Mean | Range |
| Adults | Winter19 | 28.03 | 0 - 558 | 142.78 | 0 - 2241 | 2.8 | 0 - 81 | 0.32 | 0 - 48 |
|  | Spring19 | 114.45 | 0 - 918 | 705.45 | 0 - 4653 | 0.2 | 0 - 4 | 1.01 | 0 - 36 |
|  | Summer19 | 60.09 | 0 - 864 | 343.69 | 0 - 5145 | 0 | 0 - 0 | 0.07 | 0 - 2 |
|  | Autumn19 | 56.7 | 0 - 1116 | 218.4 | 0 - 2052 | 0 | 0 - 0 | 0.79 | 0 - 72 |
|  | Winter20 | 44.79 | 0 - 354 | 140.42 | 0 - 1251 | 0 | 0 - 0 | 0.29 | 0 - 18 |
| Lambs | Summer19 | 166.35 | 0 - 792 | 5188.71 | 123 - 39942 | 0 | 0 - 0 | 154.79 | 0 - 1161 |
|  | Autumn19 | 138.93 | 0 - 1863 | 666.19 | 0 - 7917 | 0 | 0 - 0 | 3.55 | 0 - 72 |
|  | Winter20 | 137.57 | 0 - 1305 | 220.71 | 0 - 1911 | 0 | 0 - 0 | 13.04 | 0 - 108 |

**Table S2.** Model output investigating sex and season effects within age class

| Response | Term | Adults |  | Lambs |  |
| --- | --- | --- | --- | --- | --- |
|  |  | Estimate (95% CI) | pMCMC | Estimate (95% CI) | pMCMC |
| Strongyles | Intercept | 2.04 (1.77-2.34) | < 0.001 | 4.3 (3.99-4.67) | < 0.001 |
|  | SeasonSpring19 | 1.83 (1.53-2.15) | < 0.001 |  |  |
|  | SeasonSummer19 | 0.88 (0.54-1.24) | < 0.001 |  |  |
|  | SeasonAutumn19 | 0.32 (-0.03-0.69) | 0.082 | -0.3 (-0.65-0.11) | 0.148 |
|  | SeasonWinter20 | 0.42 (0.08-0.76) | 0.022 | -0.49 (-0.93--0.02) | 0.038 |
|  | SexMale | 0.97 (0.5-1.45) | < 0.001 | 0.04 (-0.34-0.4) | 0.854 |
|  | Id | 0.71 (0.41-0.98) |  | 0.19 (0-0.49) |  |
|  | units | 1.82 (1.57-2.06) |  | 2.38 (1.92-2.86) |  |
| Coccidia | Intercept | 3.97 (3.72-4.19) | < 0.001 | 8 (7.68-8.28) | < 0.001 |
|  | SeasonSpring19 | 2.13 (1.88-2.45) | < 0.001 |  |  |
|  | SeasonSummer19 | 0.98 (0.67-1.3) | < 0.001 |  |  |
|  | SeasonAutumn19 | 0.49 (0.18-0.78) | 0.004 | -2.37 (-2.67--2.05) | < 0.001 |
|  | SeasonWinter20 | 0.3 (-0.03-0.61) | 0.072 | -3.66 (-4.08--3.31) | < 0.001 |
|  | SexMale | -0.11 (-0.49-0.28) | 0.592 | 0.08 (-0.21-0.38) | 0.656 |
|  | Id | 0.4 (0.23-0.6) |  | 0.12 (0-0.31) |  |
|  | units | 1.6 (1.41-1.82) |  | 1.66 (1.34-1.96) |  |
| Capillaria | Intercept | -1.61 (-2.26--1) | < 0.001 |  |  |
|  | SeasonSpring19 | -2.75 (-3.73--1.85) | < 0.001 |  |  |
|  | SexMale | 0.63 (-0.41-1.66) | 0.264 |  |  |
|  | Id | 0.34 (0-1.32) |  |  |  |
|  | units | 5.92 (3.49-8.58) |  |  |  |
| Nematodirus | Intercept |  |  | 3.04 (2.37-3.77) | < 0.001 |
|  | SeasonAutumn19 |  |  | -5.56 (-6.58--4.61) | < 0.001 |
|  | SeasonWinter20 |  |  | -3.02 (-4--2.01) | < 0.001 |
|  | SexMale |  |  | 0.22 (-0.55-1.03) | 0.54 |
|  | Id |  |  | 0.24 (0-0.83) |  |
|  | units |  |  | 9.2 (6.95-11.55) |  |
| StrongyloidesBin | Intercept | -4.61 (-5.6--3.63) | < 0.001 | 4.3 (3.99-4.67) | < 0.001 |
|  | SeasonSpring19 | 1.88 (0.58-3.1) | 0.004 |  |  |
|  | SeasonSummer19 | -0.85 (-2.55-0.52) | 0.27 |  |  |
|  | SeasonAutumn19 | 0.86 (-0.59-2.05) | 0.204 | -0.3 (-0.65-0.11) | 0.148 |
|  | SeasonWinter20 | -1.17 (-2.88-0.26) | 0.146 | -0.49 (-0.93--0.02) | 0.038 |
|  | SexMale | 1.28 (0.21-2.38) | 0.03 | 0.04 (-0.34-0.4) | 0.854 |
|  | Id | 0.66 (0-2.06) |  | 0.19 (0-0.49) |  |
|  | units | 10 (10-10) |  | 2.38 (1.92-2.86) |  |
| MonieziaBin | Intercept | -15.57 (-17.15--14.16) | < 0.001 | 8 (7.68-8.28) | < 0.001 |
|  | SeasonSummer19 | -0.55 (-1.41-0.39) | 0.232 | -2.37 (-2.67--2.05) | < 0.001 |
|  | SeasonAutumn19 | -0.47 (-1.34-0.4) | 0.268 | -3.66 (-4.08--3.31) | < 0.001 |
|  | SexMale | 0.39 (-2.81-3.99) | 0.85 | 0.08 (-0.21-0.38) | 0.656 |
|  | Id | 51.45 (36.95-66.79) |  | 0.12 (0-0.31) |  |
|  | units | 10 (10-10) |  | 1.66 (1.34-1.96) |  |

**Table S3.** Univariate (Model Set 2) and Multivariate (Model Set 3) MCMCglmm output for Strongyle and Coccida count models from adult-restricted dataset accounting for a sex-by-season interaction.

|  | Univariate, Adults |  |  |  | Multivariate, Adults |  |  |  |
| --- | --- | --- | --- | --- | --- | --- | --- | --- |
|  | Strongyles |  | Coccidia |  | Strongyles |  | Coccidia |  |
|  | Estimate | pMCMC | Estimate | pMCMC | Estimate | pMCMC | Estimate | pMCMC |
| Intercept | 1.9 (1.6-2.17) | **< 0.001** | 4.06 (3.81-4.33) | **< 0.001** | 1.92 (1.61-2.19) | **< 0.001** | 4.06 (3.83-4.35) | **< 0.001** |
| SeasonSpring19 | 2.32 (1.99-2.67) | **< 0.001** | 2.14 (1.83-2.47) | **< 0.001** | 2.31 (1.97-2.66) | **< 0.001** | 2.13 (1.76-2.43) | **< 0.001** |
| SeasonSummer19 | 0.82 (0.48-1.19) | **< 0.001** | 0.82 (0.49-1.2) | **< 0.001** | 0.82 (0.41-1.16) | **< 0.001** | 0.81 (0.47-1.17) | **< 0.001** |
| SeasonAutumn19 | 0.53 (0.1-0.85) | **0.006** | 0.35 (0.02-0.71) | 0.052 | 0.52 (0.14-0.9) | 0.006 | 0.35 (0-0.72) | **0.048** |
| SeasonWinter20 | 0.57 (0.17-0.93) | **< 0.001** | 0.25 (-0.13-0.58) | 0.172 | 0.56 (0.17-0.94) | **< 0.001** | 0.25 (-0.14-0.58) | 0.172 |
| SexReproFemaleNoLamb | 0.66 (-0.33-1.67) | 0.2 | -0.47 (-1.3-0.36) | 0.252 | 0.62 (-0.38-1.56) | 0.236 | -0.44 (-1.21-0.43) | 0.298 |
| SexReproMale | 1.51 (0.79-2.17) | **< 0.001** | -0.52 (-1.16-0.13) | 0.126 | 1.47 (0.84-2.13) | **< 0.001** | -0.5 (-1.17-0.13) | 0.134 |
| SeasonSpring19 : SexReproFemaleNoLamb | -2.32 (-3.47--1.07) | **< 0.001** | 0.31 (-0.71-1.34) | 0.548 | -2.24 (-3.53--1.07) | **< 0.001** | 0.29 (-0.81-1.44) | 0.58 |
| SeasonSummer19 : SexReproFemaleNoLamb | -0.73 (-2.06-0.73) | 0.262 | 0.05 (-1.14-1.14) | 0.916 | -0.68 (-1.97-0.58) | 0.332 | 0.05 (-1.02-1.2) | 0.962 |
| SeasonAutumn19 : SexReproFemaleNoLamb | -0.76 (-2.03-0.56) | 0.276 | 0.25 (-0.89-1.49) | 0.672 | -0.66 (-1.99-0.67) | 0.344 | 0.28 (-0.88-1.52) | 0.674 |
| SeasonWinter20 : SexReproFemaleNoLamb | -1.25 (-2.54-0.03) | 0.06 | 0.15 (-0.98-1.34) | 0.764 | -1.2 (-2.41-0.16) | 0.076 | 0.16 (-1.16-1.23) | 0.792 |
| SeasonSpring19 : SexReproMale | -1.63 (-2.44--0.91) | **< 0.001** | 0.02 (-0.67-0.79) | 0.982 | -1.6 (-2.37--0.77) | **< 0.001** | 0.01 (-0.73-0.76) | 0.954 |
| SeasonSummer19 : SexReproMale | 0.58 (-0.3-1.48) | 0.206 | 1.05 (0.19-1.87) | **0.02** | 0.61 (-0.11-1.54) | 0.154 | 1.03 (0.17-1.89) | **0.022** |
| SeasonAutumn19 : SexReproMale | -0.91 (-1.84--0.05) | **0.046** | 0.79 (0-1.65) | 0.058 | -0.88 (-1.86-0) | 0.066 | 0.75 (-0.16-1.52) | 0.098 |
| SeasonWinter20 : SexReproMale | -0.31 (-1.25-0.52) | 0.472 | 0.32 (-0.45-1.29) | 0.452 | -0.3 (-1.2-0.64) | 0.518 | 0.3 (-0.6-1.14) | 0.468 |
| Id | 0.69 (0.44-1.03) |  | 0.4 (0.23-0.6) |  | 0.69 (0.42-0.97) |  | 0.41 (0.21-0.59) |  |
| Units | 1.69 (1.48-1.96) |  | 1.6 (1.39-1.8) |  | 1.68 (1.45-1.91) |  | 1.6 (1.4-1.81) |  |
| Strongyle : Coccidia.Id |  |  |  |  | 0.14 (-0.01-0.33) |  | 0.41 (0.21-0.59) |  |
| Strogyle : Coccida.units |  |  |  |  | 0.47 (0.32-0.62) |  | 0.47 (0.32-0.62) |  |

**Table S4.** Predicted distribution comparisons between demographic groups for seasonal changes

| Seasonal Change | Comparison Levels | Strongyles |  |  |  |  | Coccidia |  |  |  |  |
| --- | --- | --- | --- | --- | --- | --- | --- | --- | --- | --- | --- |
|  |  | Mean | Lower | Upper | P | Sig | Mean | CI Lower | CI Upper | P | Sig |
| Spring19-Winter19 | FemaleNoLamb-FemaleLamb | -2.28 | -3.41 | -1.14 | <0.001 | *** | 0.3 | -0.8 | 1.34 | 0.57 |  |
|  | Male-FemaleLamb | -1.65 | -2.47 | -0.96 | <0.001 | *** | 0.02 | -0.7 | 0.8 | 0.99 |  |
|  | FemaleLamb-FemaleNoLamb | 2.28 | 1.14 | 3.41 | <0.001 | *** | -0.3 | -1.34 | 0.8 | 0.57 |  |
|  | Male-FemaleNoLamb | 0.64 | -0.71 | 1.85 | 0.35 |  | -0.28 | -1.48 | 1.07 | 0.64 |  |
|  | FemaleLamb-Male | 1.65 | 0.96 | 2.47 | <0.001 | *** | -0.02 | -0.8 | 0.7 | 0.99 |  |
|  | FemaleNoLamb-Male | -0.64 | -1.85 | 0.71 | 0.35 |  | 0.28 | -1.07 | 1.48 | 0.64 |  |
| Summer19-Spring19 | FemaleNoLamb-FemaleLamb | 1.6 | 0.46 | 2.84 | 0.01 | ** | -0.26 | -1.29 | 0.9 | 0.63 |  |
|  | Male-FemaleLamb | 2.24 | 1.47 | 3.09 | <0.001 | *** | 1.04 | 0.17 | 1.81 | 0.01 | * |
|  | FemaleLamb-FemaleNoLamb | -1.6 | -2.84 | -0.46 | 0.01 | ** | 0.26 | -0.9 | 1.29 | 0.63 |  |
|  | Male-FemaleNoLamb | 0.64 | -0.58 | 2.11 | 0.36 |  | 1.29 | 0.04 | 2.54 | 0.04 | * |
|  | FemaleLamb-Male | -2.24 | -3.09 | -1.47 | <0.001 | *** | -1.04 | -1.81 | -0.17 | 0.01 | * |
|  | FemaleNoLamb-Male | -0.64 | -2.11 | 0.58 | 0.36 |  | -1.29 | -2.54 | -0.04 | 0.04 | * |
| Autumn19-Summer19 | FemaleNoLamb-FemaleLamb | -0.05 | -1.48 | 1.18 | 0.94 |  | 0.24 | -0.91 | 1.4 | 0.65 |  |
|  | Male-FemaleLamb | -1.51 | -2.44 | -0.63 | <0.001 | *** | -0.25 | -1.09 | 0.57 | 0.56 |  |
|  | FemaleLamb-FemaleNoLamb | 0.05 | -1.18 | 1.48 | 0.94 |  | -0.24 | -1.4 | 0.91 | 0.65 |  |
|  | Male-FemaleNoLamb | -1.46 | -3.15 | -0.08 | 0.06 |  | -0.5 | -1.85 | 0.84 | 0.44 |  |
|  | FemaleLamb-Male | 1.51 | 0.63 | 2.44 | <0.001 | ** | 0.25 | -0.57 | 1.09 | 0.56 |  |
|  | FemaleNoLamb-Male | 1.46 | 0.08 | 3.15 | 0.06 |  | 0.5 | -0.84 | 1.85 | 0.44 |  |
| Winter20-Autumn19 | FemaleNoLamb-FemaleLamb | -0.46 | -1.82 | 0.8 | 0.47 |  | -0.14 | -1.35 | 1.04 | 0.8 |  |
|  | Male-FemaleLamb | 0.62 | -0.32 | 1.61 | 0.22 |  | -0.48 | -1.29 | 0.41 | 0.31 |  |
|  | FemaleLamb-FemaleNoLamb | 0.46 | -0.8 | 1.82 | 0.47 |  | 0.14 | -1.04 | 1.35 | 0.8 |  |
|  | Male-FemaleNoLamb | 1.08 | -0.4 | 2.54 | 0.15 |  | -0.34 | -1.75 | 0.97 | 0.63 |  |
|  | FemaleLamb-Male | -0.62 | -1.61 | 0.32 | 0.22 |  | 0.48 | -0.41 | 1.29 | 0.31 |  |
|  | FemaleNoLamb-Male | -1.08 | -2.54 | 0.4 | 0.15 |  | 0.34 | -0.97 | 1.75 | 0.63 |  |

